# Asleep at the Wheel: Forward Genetic ENU Mutagenesis Screen for Mouse Models of Chronic Fatigue Identifies a Mutation in *Slc2a4* (GLUT4)

**DOI:** 10.1101/163378

**Authors:** Marleen H. M. de Groot, Carlos M. Castorena, Vivek Kumar, Jennifer A. Mohawk, Newaz I. Ahmed, Joseph S. Takahashi

**Author notes:** Present Address: The Jackson Laboratory, Bar Harbor, ME 04609. To whom correspondence should be addressed: Dr. Joseph S. Takahashi, HHMI, Department of Neuroscience, University of Texas Southwestern Medical Center, 5323 Harry Hines Blvd., NA4.118, Dallas, TX 75390-9111 USA. Classification: Biological Sciences – Genetics.

## Abstract

In a screen of voluntary wheel-running behavior designed to identify genetic mouse models of chronic fatigue in ENU mutagenized C57BL/6J mice, we discovered two lines that showed aberrant wheel-running patterns. These lines both stem from a single original founder identified as a low body-weight candidate in a recessive screen. Progeny from both of these lines showed the abnormal wheel-running behavior, with affected mice showing significantly lower daily activity levels than unaffected mice. They also exhibited low amplitude circadian rhythms, consisting of lower activity levels during the normal active phase, and increased levels of activity during the rest or light phase, but only a modest alteration in free-running period. Their activity is not consolidated into longer bouts, but is frequently interrupted with periods of inactivity throughout the dark phase of the light-dark (LD) cycle. As seen with the low body weight, expression of the behavioral phenotypes in offspring of strategic crosses was consistent with a recessive heritance pattern. Mapping of these phenotypic abnormalities showed linkage to a single locus on chromosome 11, and whole exome sequencing (WES) identified a single point mutation in the *Slc2a4* gene encoding the GLUT4 insulin-responsive glucose transporter. The single nucleotide change (A to T) was found in the distal end of exon 10, and results in a premature stop (Y440*). To our knowledge, this is the first time a mutation in this gene has been shown to result in extensive changes in general behavioral patterns.

**SIGNIFICANCE STATEMENT:** Chronic fatigue is a debilitating and devastating disorder with widespread consequences for both the patient and the persons around them, but effective treatment strategies are lacking. The identification of novel genetic mouse models of chronic fatigue may prove invaluable for the study of its underlying physiological mechanisms and for the testing of treatments and interventions. A novel mutation in *Slc2a4* (GLUT4) was identified in a forward mutagenesis screen because affected mice showed abnormal daily patterns and levels of wheel running consistent with chronic fatigue. This new mouse model may shed light on the pathophysiology of chronic fatigue.

## INTRODUCTION

Fatigue is a debilitating symptom comorbid with many disorders and disease states, and may also present without readily identifiable associated disorders, the latter referred to as idiopathic fatigue (1). Fatigue persisting 6 months or more is considered chronic fatigue and if additional criteria are met, a diagnosis of Chronic Fatigue Syndrome (CFS) can be made (2-4). The Centers for Disease Control and Prevention define chronic fatigue syndrome as “a debilitating and complex disorder characterized by intense fatigue that is not improved by bed rest and that may be worsened by physical activity or mental exertion”. Fatigue is associated with increased disability, impaired functioning and increased mortality (5-7). Its negative impact on activity and overall quality of life further increases the need for adequate and effective treatment, although this has remained elusive.

The development of appropriate animal models to study the underlying mechanisms of fatigue is crucial, but is hampered by the lack of a widely accepted operational definition of fatigue (8, 9). Currently there are several different types of mouse models, including those based on forced activity (e.g., treadmill or swimming), neurodegenerative disorders (e.g., Parkinson’s or Huntington’s disease), cancer and its treatments, or aging, but by far the most common types of models involve fatigue following an immune challenge (e.g., those mimicking viral or parasitic infection) (10-16). A commonality shared by all of these model systems is a resulting alteration of behavioral output, such as voluntary wheel running (9).

The mouse is an ideal model organism to investigate physiological and neurological pathways involved in fatigue (9, 17). Although the psychological and emotional aspects of fatigue may be somewhat difficult to study in mice, the behavioral components are more readily comparable. Models based on reduced or altered voluntary wheel-running activity mimic the reduced voluntary physical activity exhibited by humans with fatigue. Wheel running is a robust behavior in rodents, and is amenable to quantitative analysis (17-19). Some characteristics that may imply fatigue in mice when tested on running wheels include reduced daily amount of activity, changes in pattern or day-night distribution, a lack of consolidated running with an increase in the number of distinct bouts, or a delayed onset or shortened duration of the daily active phase.

There is some indication that there are genetic components related to the development and manifestation of fatigue (20-23). Polymorphisms in several human candidate genes (e.g., the adrenergic receptor a1 (ADRA1A), the serotonin transporter (5-HTT or SLC6A4) or receptor (HTR2A), tyrosine hydroxylase (TH), corticosteroid-binding globulin (CBG), corticotropin releasing hormone receptor 1 (CRHR1), the cytokine IL-1B, neuronal PAS domain protein 2 (NPAS2), the nuclear receptor subfamily 3; group C, member 1 glucocorticoid receptor (NR3C1), and the glutamate receptor - ionotropic kinase 2 (GRIK2)) have been linked to both the occurrence and severity of fatigue symptomology (24-30). A study comparing chronic fatigue syndrome patients to healthy controls using a single nucleotide polymorphism (SNP) chip with over 900,000 SNPs identified 442 candidates associated with chronic fatigue syndrome, and highlighted two genes never before associated with fatigue (C-type lectin domain family 4, member M (CLEC4M) and coiled-coil domain containing 157 protein (CCDC157)) (31). In addition, mice with mutations in several genes have shown phenotypes consistent with fatigue (e.g., corticosteroid-binding globulin (CBG), recombinase activating gene 2 (RAG2), and interleukin-10 (IL-10)) (32-34).

We know that random mutagenesis paired with forward genetic screening has proven invaluable in establishing and expanding understanding of gene function (35-43). In the field of circadian rhythms, forward genetics has been a powerful tool for identifying both clock and clock-controlled genes (44). We systematically screen randomly mutagenized mice on running wheels in search of lines exhibiting alterations in voluntary wheel-running activity. We believe that the identification of mouse lines that harbor mutations resulting in altered behavioral patterns consistent with chronic fatigue, and the subsequent identification of the responsible gene(s) or pathways, will contribute significantly to our understanding of the pathophysiology of this debilitating disorder.

As part of a larger phenotype-driven ENU mouse mutagenesis project (45), we screened mice for abnormalities in the expression of voluntary wheel-running activity, with the hope of identifying genes involved in behavioral chronic fatigue. We describe the identification of two independent lines of mice stemming from a single mutagenized founder showing profound alterations in amount, pattern and distribution of daily wheel running resulting from a single point mutation (*twiggy*; MGI:5805978) in *Slc2a4* encoding the insulin-responsive glucose transporter, GLUT4. To our knowledge, this is the first description of impaired functioning of this gene resulting in extensive behavioral abnormalities. The finding that a mutation in this gene results in widespread behavioral modifications and altered expression of daily activity patterns indicates that its role is likely more diverse than has previously been appreciated.

## RESULTS

### Mice show abnormal, low amplitude voluntary wheel-running activity patterns

In a voluntary wheel-running screen designed to isolate mice with heritable abnormal patterns of behavior, we identified two lines of mice showing abnormal activity consistent with chronic fatigue (Fig. 1). These two lines both stemmed from a single, third generation (G3) founder female identified as having low body weight (Fig. S1 A), which we named ‘*twiggy*’ and which was backcrossed to wildtype C57BL/6J to produce N2 progeny (Fig. S1 B). Two additional G3 pedigrees stemming from the same mutagenized founder (G0) male also contained affected G3 mice with low body weight (2 out of 8 mice and 3 out of 26 mice, respectively) but none of these G3s produced enough offspring for heritability testing. N2 progeny of the *twiggy* G3 founder female showed body weights within the normal range, but an intercross of N2 mice produced a subset (~5-8 out of 57) that showed the low body-weight phenotype (Fig. S1B). A subsequent backcross of affected N2F2 mice to wildtype C57BL/6J produced N3 progeny that were unaffected, indicating that this phenotype is expressed in a non-sex-linked, recessive heritance pattern. Despite the low body-weight phenotype and overall smaller appearance, affected animals were completely healthy, and there was no observed effect on reproductive fitness.

**Fig. 1.**
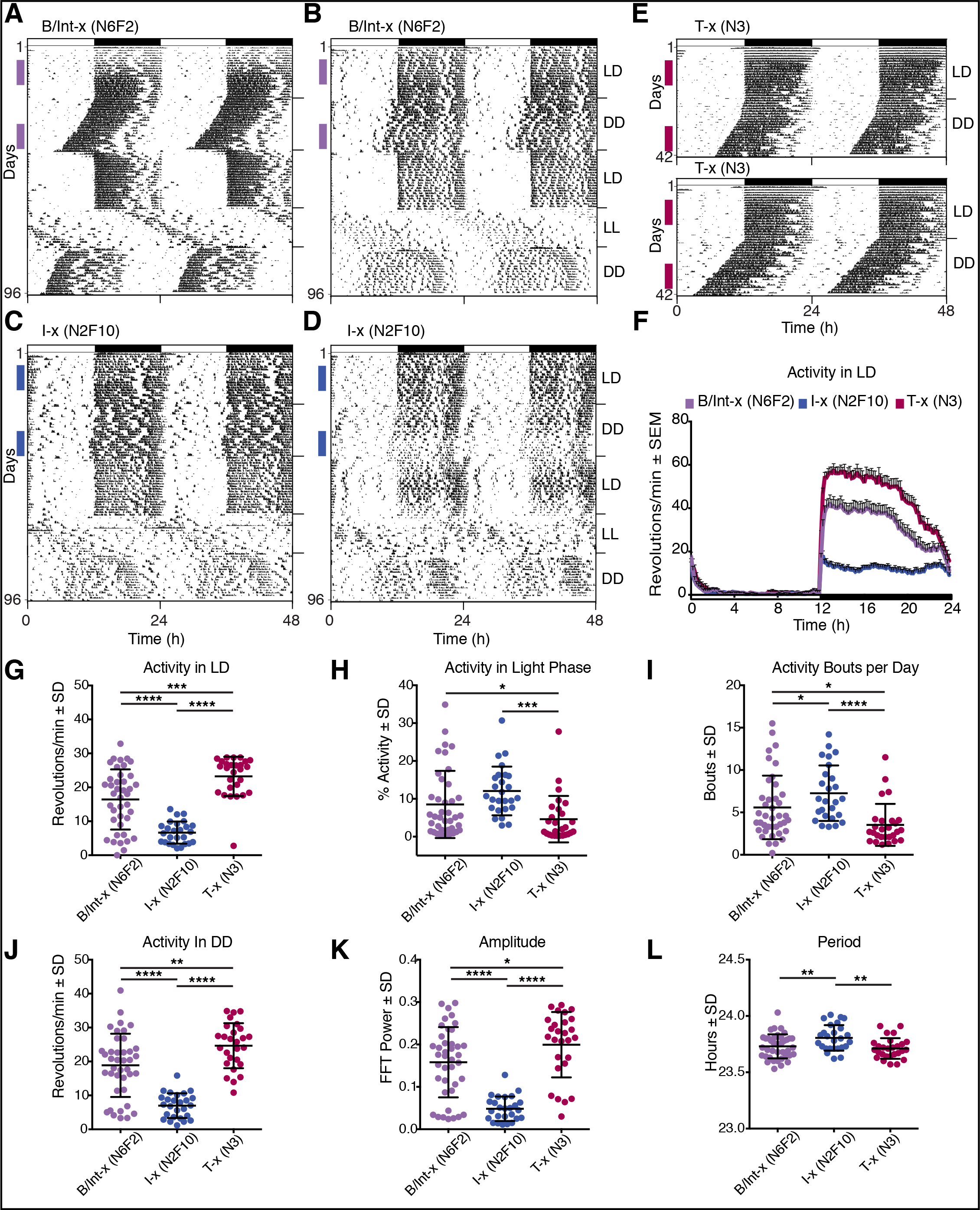
Mice from two lines show abnormal, low amplitude voluntary wheel-running activity patterns. (A) Representative actogram of a B/Int-x (N6F2) mouse showing a normal circadian activity pattern, double plotted for ease of visualization, with each horizontal line representing 48h and successive days (1-96) plotted along the Y-axis. Wheel revolutions are depicted as black marks on the horizontal lines. Mice were initially recorded under a 12h light:12h dark (LD) cycle shown in the bar across the top of the graph (black = dark phase). After 3 weeks, mice were recorded in constant darkness (DD) for 3 weeks, in 12:12 LD for 3 weeks, in constant light (LL) for 2 weeks and a final 2 weeks in DD (indicated on right Y-axis). Wheel running activity shows a normal nocturnal pattern during recording under LD with high levels of activity concentrated in the dark phase and low activity during the light phase. In DD, the rhythm free-runs with a period of ~23.7h, re-entrains normally upon reintroduction of the LD cycle, and free-runs with a lengthened period under LL. Colored vertical bars on the left side of the actogram indicate days used for quantification of wheel-running behavior. Representative actogram of a B/Int-x (N6F2) (B) and two I-x (N2F10) (C&D) mice showing an abnormal circadian activity pattern with plotting conventions and recording conditions as described for (A). (E) Representative actograms of two T-x (N3) mice recorded for 3 weeks in LD followed by 3 weeks in DD. Plotting conventions are as described for (A). (F) Average waveforms for activity recorded in LD for mice in the B/IntX (N6F2; n=40), I-x (N2F10; n=26), and T-x (N3; n=27) groups. Time (24h) is plotted along the X-axis and average number of wheel revolutions (± SEM) is plotted along the Y-axis. The LD cycle is shown in the bar across the bottom of the graph with lights off from 12 to 24 indicated in black. Average number of wheel revolutions per min over 24h (G), % of daily activity occurring in the light phase (H), and number of discreet bouts of activity occurring each day (I) during recording under LD for mice in the B/IntX (N6F2; n=40), I-x (N2F10; n=26), and T-x (N3; n=27) groups are plotted. Average number of wheel revolutions per min over 24h (J), amplitude (K) and period (L) of the wheel-running rhythm recorded in DD for these same mice is also shown. Means ± SD with individual data points are plotted in G-L with significant differences among groups indicated (* = p < 0.05, ** = p < 0.01, *** = p < 0.001, **** = p < 0.0001).

Mice were maintained in two distinct lines following initial heritability testing (Fig. S1 C and D; Table S1). The first is a backcross/intercross (B/Int-x) line generated and maintained by backcrossing affected (presumably homozygous) mice to C57BL/6J, and then intercrossing the resulting offspring, producing alternating generations of obligate heterozygous carrier “B/Int-x (N#)” mice, and “B/Int-x (N#F2)” mice of all 3 possible genotypes (+/+, +/m, and m/m; ~25%, ~50% and ~25%, respectively; Fig. S1 C and D – left panels; Table S1; where N# denotes the number of backcross generations). The second is an incross (I-x) line generated and maintained by crossing affected siblings with each other over successive generations beginning with the first N2F2 progeny shown in Fig. S1 B, to produce a homozygous line (Fig. S1 C and D – right panels). The body weights of 8 sequential generations of each line are shown in Fig. S1 C (males) and D (females), and, while all 8 I-x generations show body weights lower than the screening mean, mice in the B/Int-x show a distinct generation-alternating pattern of results.

Because of the low body weight, we measured fasting blood glucose from individuals of both the B/Int-x and I-x lines. Although body weights following a 4h fast were significantly lower for both males and females of the I-x line than those of the B/Int-x line (Fig. S1 E), there was no significant difference between the two lines in fasting blood glucose (Fig. S1 F). Values for all mice tested were within the normal blood glucose range for C57BL/6J mice reported by Jackson Labs (Center for Genome Dynamics; CGDpheno1 glucose (plasma GLU, 4h fast); phenome.jax.org).

When mice stemming from these lines were housed in cages with free access to a running wheel, a number of mice in the B/Int-x (N6F2) line showed a hypoactive and disrupted pattern of voluntary wheel-running activity (Fig. 1 A and B). Representative activity plots (actograms) of two mice of the B/Int-x (N6F2) line are shown, with one exhibiting normal levels and patterns of activity (Fig. 1 A) and one showing the abnormal phenotype (Fig. 1 B). In contrast, all the mice in the I-x (N2F10) line showed this same aberrant pattern (Fig. 1 C and D), consisting of low activity levels, increased bouts of activity during the light or rest phase, non-consolidated or disrupted nocturnal activity, and irregular onsets of the main nocturnal bout. When affected mice from the I-x (N2F10) line were backcrossed to C57BL/6J mice in a test cross (T-x (N3); Table S1), none of the progeny showed the phenotype, and circadian, or daily, wheel-running behavior was indistinguishable from that of wildtype mice (Fig. 1 E). Average waveforms of 10 days of recording under a light-dark (LD) cycle clearly show that overall wheel-running activity levels are significantly decreased in mice of the I-x (N2F10) line (Fig. 1 F). The B/Int-x (N6F2) line included some mice that showed the phenotype and some that didn’t, whereas none of the T-x (N3) mice were affected, and these showed a normal circadian pattern of activity with high levels of nocturnal wheel running.

Quantification of total activity (Fig. 1 G), light phase activity (Fig. 1 H), and number of activity bouts per day (Fig. 1 I) recorded during the LD cycle, showed significant differences among progeny of the three types of crosses, with mice of the I-x (N2F10) line showing low activity, with higher levels of activity during the rest phase and higher numbers of distinct activity bouts. Differences in activity levels were maintained when mice were recorded in constant darkness (DD; Fig. 1 J), and the amplitude of the wheel running rhythm in affected mice was significantly lower than in unaffected mice (Fig. 1 K). The period of the free-running circadian rhythm recorded in DD was significantly longer for mice of the I-x (N2F10) line (Fig. 1 L), although these values fall within the normal range for wildtype C57BL/6J (18).

### Frequent rest bouts interrupt nocturnal wheel-running activity in affected mice

In order to characterize further the abnormal behavioral pattern these mice exhibit, we assessed their behavioral profiles during the dark phase using both time-lapsed photography (Fig. 2) and video recording (Fig. 3). Images captured once every min for 15h beginning at the start of the dark phase for 3 individual wildtype C57BL/6J (Fig. S2) and 5 individual I-X (N2F14) mice (Fig. S3) were scored on a range of 9 behavioral states. The I-X (N2F14) mice were each recorded twice, with one week occurring between recording sessions (Fig. S3). The ethogram of scored behaviors for the representative wildtype C57BL/6J mouse shows extended bouts of wheel running with short interruptions for drinking, eating, and general cage activity (Fig. 2 A). This mouse only adopted a body posture consistent with sleep during the end of the end of the dark phase (i.e., after 3:00 h). The amount of time spent in the various behaviors for the first 6h of the dark phase (18:00 – 0:00 h) are represented in a pie chart for this individual mouse in Fig. 2 C and averaged for all 3 wildtype mice in Fig. 2 D. The pattern of behaviors shown by the representative I-x (N2F14) mouse (Fig. 2 B) does not resemble that recorded for wildtype mice. Consolidated, extended bouts of wheel running are absent, and running is interrupted with periods of time when the mouse is near the wheel, but is not active. During these periods of time, the mouse is seen to adopt a low, crouched posture indicative of sleep (Fig. 2 G – right panel). The distribution of behaviors for the first 6h of the dark phase (18:00 – 0:00 h) for this mouse is shown in Fig. 2 E, and for all 5 I-x (N2F14/15) mice recorded twice in Fig. 2 F. Most notably, running is significantly decreased for these mice compared to wildtype (Fig. 2 C-F), Even more surprising, all 5 I-X (N2F14/15) mice show sleep postures during this 6h time period on both recording sessions, whereas wildtype mice were never observed to do this (Fig. S2 and S3).

**Fig. 2.**
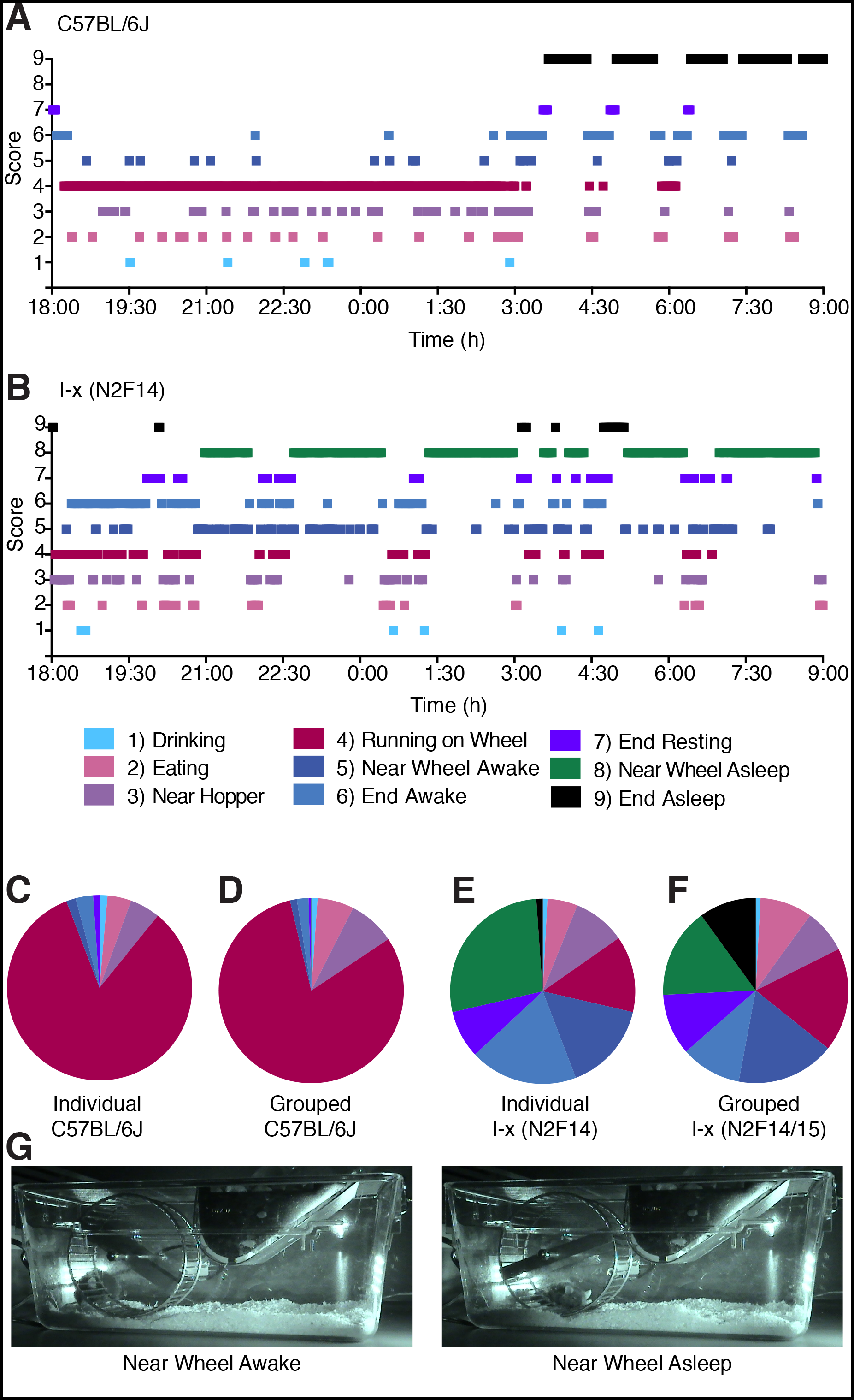
Nocturnal wheel-running activity is interrupted by frequent rest bouts in affected mice. Ethogram depicting behaviors, states or locations observed in images captured once every min for 15h beginning at lights off for a wildtype C57BL/6J (A) and an I-x (N2F14) (B) mouse, with time plotted along the X-axis. Each photograph was scored on a 9 item scale using the following designations: 1) drinking, 2) eating, 3) near hopper, 4) running on wheel, 5) near wheel awake, 6) end awake, 7) end resting, 8) near wheel asleep, and 9) end asleep, where ‘end’ refers to the end of the cage furthest away from the wheel. Color code for each score is shown in the legend below (B). Pie charts showing the amount of time spent in each of the 9 behavioral states for the first 6h of recording for the individual plotted in A (C), and averaged for all 3 wildtype C57BL/6J mice (D), the individual plotted in B (E), and averaged for all 10 I-x (N2F14/15) mice (F) are shown. Note: 5 individual I-x (N2F14/15) mice were recorded twice each with ~one week between recording days. Pictures in (G) show a representative mouse scored as “near wheel awake” (right) and “near wheel asleep” (left). See also data for individual mice plotted in Fig. S2 and S3.

**Fig. 3.**
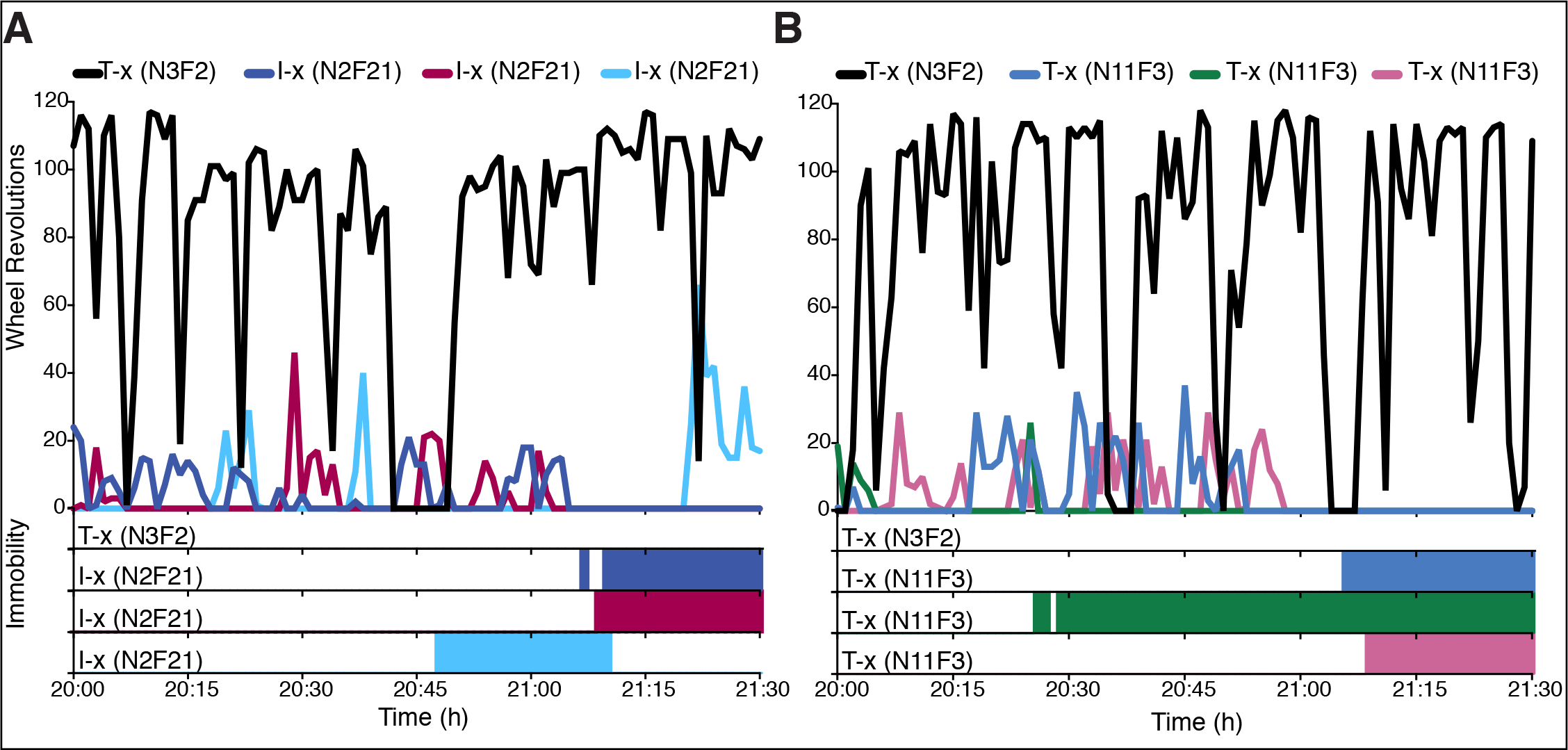
Video recording shows bouts of sleep during active phase in affected mice. Total number of wheel revolutions recorded for 2h to 3.5h after lights off for a T-x (N3F2) mouse and 3 individual I-x (N2F21) mice (A), and a different T-x (N3F2) and 3 individual T-x (N11F3) mice (B) are plotted, with periods of prolonged immobility consistent with sleep indicated in colored bars along the X-axis below. Periods > 40s of total immobility with low posture were scored as sleep.

Videos recorded during the first few hours of the dark phase for three individual mice of the I-x (N2F21) and T-x (N11F3) generations were scored and the data are plotted along with those for two unaffected T-x (N3F2) mice (Fig. 3 A and B). Mice of both affected groups showed extremely low wheel-running levels compared to T-x (N3F2) unaffected mice (Fig. 3 A and B – upper panels). The T-x (N3F2) mice never showed behavioral arrest and lowered posture consistent with sleep during this time of the day, whereas all three individuals of the other two lines did (Fig. 3 A and B – lower panels).

### Abnormal phenotypes map to a single locus on chromosome 11

To map the mutation, we crossed affected mice from both the I-x and the B/Int-x lines to C57BL/10J mice, and then intercrossed the resulting M-x (F1) progeny (n=92) to produce M-x (F2) mice (Table S1). In total, 207 M-x (F2) mice from 9 independent matings were produced and were tested on running wheels under LD and DD conditions for the presence of the behavioral phenotype (Fig. 4). All data were compared to F1 and F2 mice generated from crosses of wildtype C57BL/6J and C57BL/10J mice (designated as WTB6B10F1 (n=63) and WTB6B10F2 (n=112), respectively). Representative actograms recorded from two M-x (F2) mice showing the low-amplitude, low activity, highly disrupted behavioral wheel-running phenotype are shown in Fig. 4 B and C, with an unaffected mouse shown in Fig. 4 A. No WTB6B10F1, WTB6B10F2 or M-x (F1) mouse exhibited this constellation of behavioral phenotypes, although some individuals did show some of its component characteristics (e.g., low activity but no increased running in the light phase or no lack of consolidation of nocturnal activity). Mice of the M-x (F2) generation showed decreased activity recorded under LD (Fig. 4D) and DD (Fig. 4E), increased amount of activity in the light phase (Fig. S4 A), and a lower amplitude rhythm recorded in DD (Fig. 4F). There was no significant difference found in number of activity bouts per day (Fig. S4 B), and the period of the free-running rhythm was not altered for M-x (F2) mice, although interestingly, the WTB6B10F2 mice did show a significantly shorter period (Fig. S4 C).

**Fig. 4.**
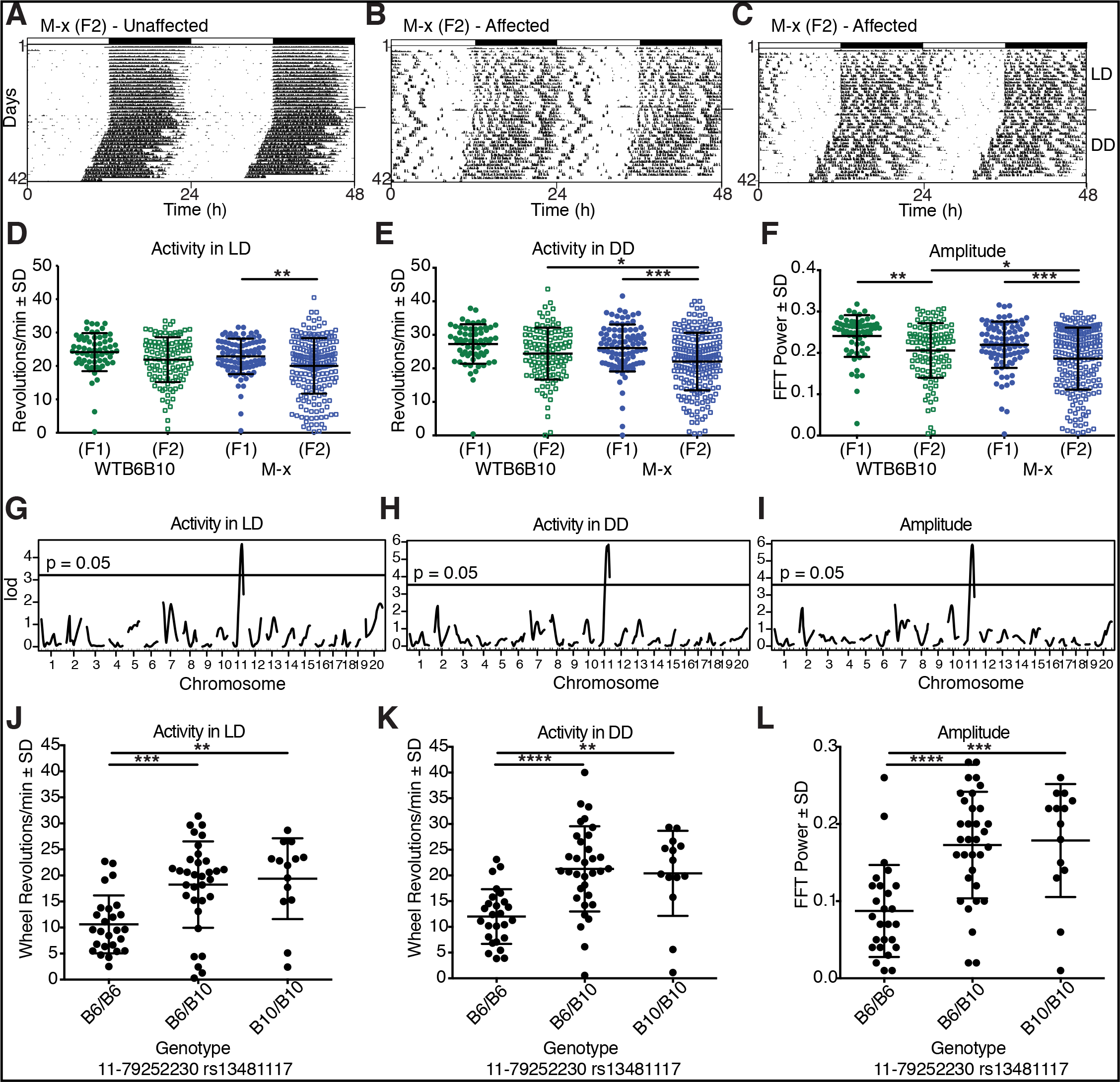
Abnormal phenotypes map to locus on chromosome 11. Representative actograms of an unaffected (A) and two affected (B & C) M-x (F2) mice, which were recorded for 3 weeks in LD followed by 3 weeks in DD. Plotting conventions are as those described for Fig. 1A. Average number of wheel revolutions per min recorded in LD (D) or DD (E) and amplitude of the free-running rhythm in DD (F) for mice in the WTB6B10 (F1; n=63), WTB6B10 (F2; n=112), M-x (F1; n=92) and M-x (F2; n=207) groups are shown. Genome wide linkage plots for average number of wheel revolutions per min recorded in LD (G) or DD (H) and amplitude of the free-running rhythm in DD (I) all show linkage to a locus on chromosome 11, with chromosomal location plotted along the X-axis. Average number of wheel revolutions per min recorded in LD (J), or DD (K) and amplitude of the free-running rhythm in DD (L) for mice with B6/B6 (n=26), B6/B10 (n=33), or B10/B10 (n=14) genotypes at a SNP marker (rs13481117) located on chromosome 11 at position 79252230 are plotted. Means ± SD and individual data points are plotted and significant differences among groups are indicated (* = p < 0.05, ** = p < 0.01, *** = p < 0.001, **** = p < 0.0001). See Fig. S4 for related data.

Using a genome-wide SNP panel of markers designed to differentiate C57BL/6J from C57BL/10J (Table S2), the genotypes of 74 selected affected and unaffected M-x (F2) mice were determined, and were assessed for linkage to the various phenotypic measurements. Significant log odds ratio (LOD) peaks with linkage to a single locus on chromosome 11 were found for the amount of activity recorded in LD (Fig. 4 G) and DD (Fig. 4 H), and amplitude of the rhythm in DD (Fig. 4 I). No significant linkage was found for the percentage of activity occurring during the light phase (Fig. S4 D and G), the number of activity bouts occurring per day (Fig. S4 E and H), or the free-running period in DD (Fig. S4 F and I). Activity levels in LD (Fig. 4 J), and in DD (Fig. 4 K), as well as the amplitude of the rhythm in DD (Fig. 4 L), are plotted as a function of genotype at SNP marker rs13481117 located on chromosome 11, position 79252230. Mice homozygous for the C57BL/6J allele (designated as “B6/B6”) show lower activity levels in both LD and DD with a lower amplitude circadian rhythm.

### A point mutation in *Slc2a4* encoding GLUT4 is identified in affected mice

In order to identify the gene responsible for the abnormal phenotypes described, we performed whole-exome sequencing (WES) of pooled DNA collected from mice of the I-x (N2F20) generation, and those from a test cross stemming from affected B/Int-x (N11F2) mice (T-x (N11F3); N=16 for each line). Representative activity records of mice in the various generations used for this experiment are shown in Fig. S5 A-F. Body-weight data collected at 8 weeks of age for both male and female mice in the I-x (N2F20) and T-x (N11F3) generations, as well as for those of the B/Int-x (N11) and B/Int-x (N1F2) used to produce the T-x (N11F3) mice, reveal that mice of both lines chosen for WES have significantly lower body weight (Fig. S5 G). In addition, mice of both presumptive homozygous lines (I-x (N2F20) and T-x (N11F3)) showed significantly lower levels of activity in LD compared to mice from the B/Int-x (F1) and B/Int-x (F2) generations (Fig. S5 H). Importantly, activity levels and wheel running patterns did not differ between these two presumptive homozygous lines, indicating that both were, in fact, homozygous for the same genetic mutation.

WES results revealed a single point mutation in the *Slc2a4* gene encoding GLUT4, the insulin-responsive glucose transporter (Fig. 5). No other gene within the linked region of chromosome 11 had polymorphisms represented in both of the samples analyzed. Remarkably, all reads covering this region of the genome for both of the pooled samples showed this same SNP (Fig. 5 A – red arrows). For clarity, 19 representative reads from each of the pooled samples are shown at higher magnification (Fig. 5 A – lower panel). A single nucleotide substitution (A to T) within the distal end of exon 10 of the gene results in a premature stop (Y440*; see Fig. 5 B). Genotyping for this SNP revealed that all mice in the 2 test cross lines were homozygous for the mutant allele of this gene, whereas mice in the B/Int-x lines showed the expected Mendelian ratio of genotypes at this mutant locus. Surprisingly, western blot analysis showed that homozygous mice had higher levels of protein in the gastrocnemius muscle than heterozygotes and wildtype littermates (Fig. 5 C), indicating that the mutation appears to result in protein accumulation or increased expression.

**Fig. 5.**
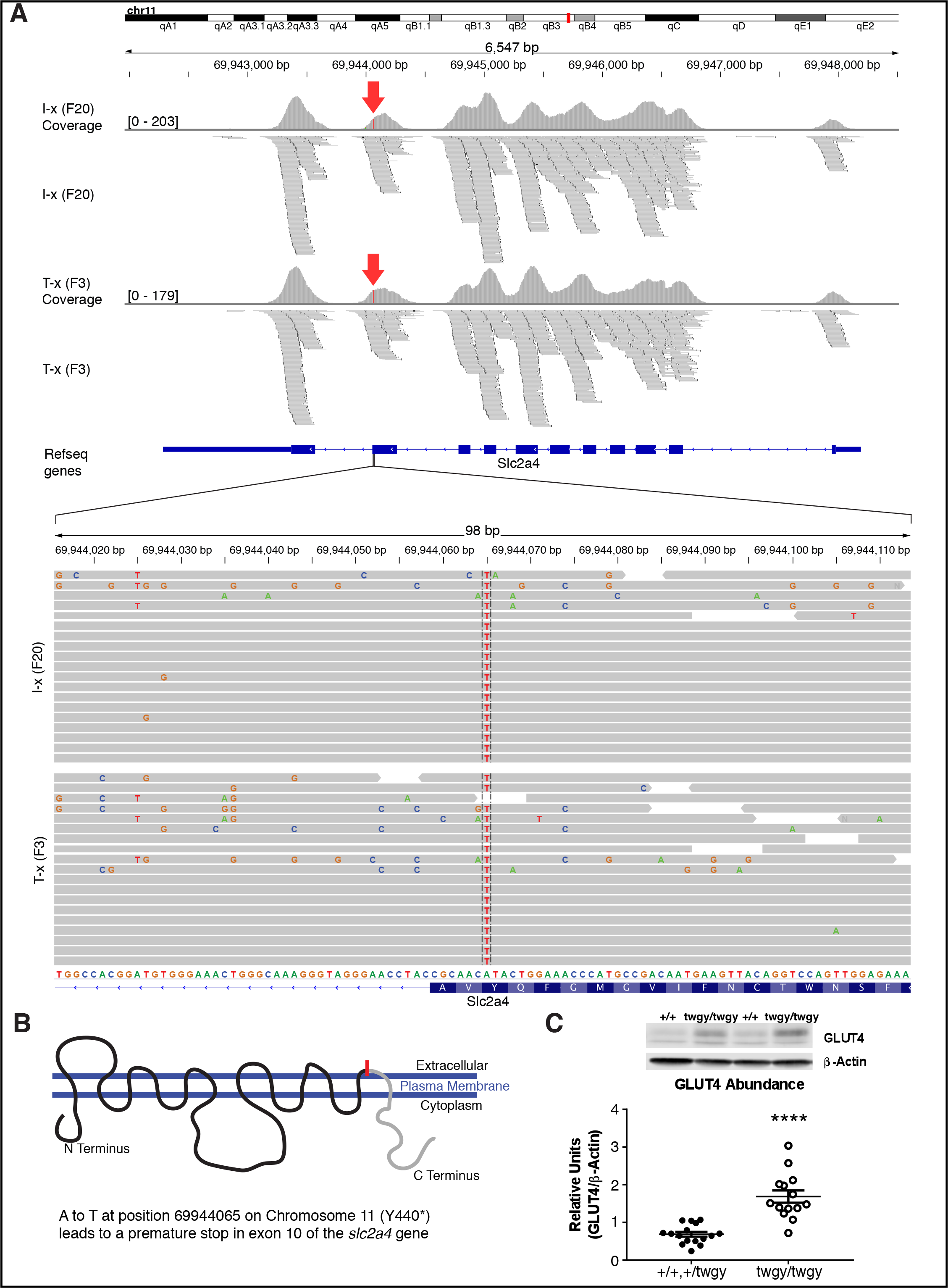
Affected mice harbor a single point mutation in *Slc2a4* encoding the GLUT4 insulin-responsive glucose transporter. Results of whole-exome sequencing of spleen genomic DNA reveal a single amino acid change (A to T) at position 69944065 within the *Slc2a4* gene on chromosome 11 (A), indicated with a red arrow on the coverage plots. All reads at this locus for the I-x (N2F20) and the T-x (N11F3) samples (n=16 per group, pooled) showed this single nucleotide alteration (indicated by red line), and 19 representative reads from each sample are shown at higher magnification at the bottom of the plot. A cartoon of the GLUT4 insulin-responsive glucose transporter with its 12 trans-membrane domains is shown in (B) with the A to T SNP location and resulting premature stop (Y440*) in exon 10 indicated with a red line. The mutation falls within the last extracellular loop and likely leads to a truncation of the cytoplasmic C-Terminus domain (indicated in grey). Protein abundance of GLUT4 in the gastrocnemius muscle of representative wildtype (+/+) and homozygous mutant (twgy/twgy) mice is shown in C with means ± SD and individual data points of +/+ or +/twgy (n=16) and twgy/twgy (n=14) plotted below. **** indicates significant difference between genotypes (p < 0.0001). See Fig. S5 for related data.

Despite the higher levels of accumulated protein, mice homozygous for the *twiggy* mutation show phenotypes similar to those shown by GLUT4 knockout mice (46-50). Despite normal fasting glucose levels (Fig. 6 B and also Fig. S1 F), *Slc2a4^twgy/twgy^* mice had impaired glucose clearance with significantly elevated blood glucose levels 15, 30 and 45 min after glucose injection (Fig. 6 A). Both +/twgy and twgy/twgy male mice showed decreased body weight, although at this age female body weights did not differ significantly among genotypes (Fig. 6 C). Like GLUT4 knockout mice (46), twgy/twgy of both sexes had enlarged hearts (Fig.6 D). The *twiggy* mutation in *Slc2a4* resulted in impaired exercise tolerance, with twgy/twgy mice showing shorter times (Fig. 6 E) and distance (Fig. 6 F) to exhaustion upon testing on a treadmill. Homozygous twgy/twgy mice also had elevated blood glucose (Fig. 6 G) and decreased lactate (Fig. 6 H) levels following completion of the treadmill task.

**Fig. 6.**
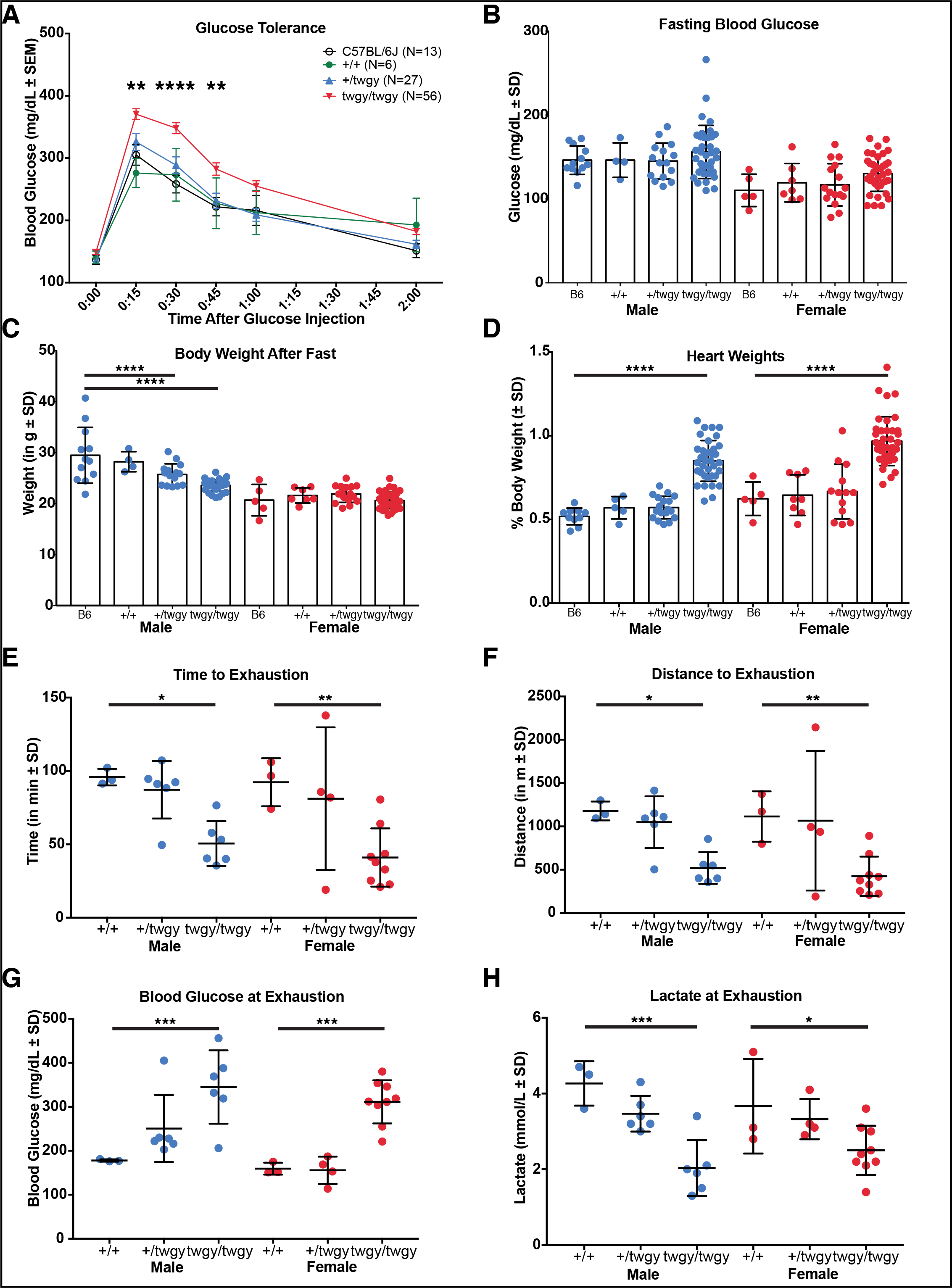
The *twiggy* mutation of *Slc2a4* results in impaired glucose tolerance, increased heart weight, low body weight, and decreased exercise tolerance. (A) Glucose tolerance response for C57BL/6J (n=13), wildtype (+/+; n=6), heterozygous (+/twgy; n=27) and homozygous (twgy/twgy; n=56) mice. Blood glucose (mg/dL ± SEM) is plotted on the Y-axis with time since I.P. injection of glucose plotted along the X-axis. Significant differences between twgy/twgy and C57BL/6J at 15, 30 and 45 min is indicated (** = p < 0.01, **** = p < 0.0001). +/+ and +/twgy did not differ from C57BL/6J at any time point. (B) Blood glucose (mg/dL ± SD) following a 4h fast in the middle of the light phase for male (blue, left) and female (red, right) C57BL/6J (B6; n=12 M & n=5 F), +/+ (n=4 M & n=7 F), +/twgy (n=15 M & n=16 F) and twgy/twgy (n=36 M & n=3 8F) is plotted. (C) Body weights of the same mice shown in B taken after a 4h fast. Heart weights (as a percentage of total body weight) of male (blue, left) and female (red, right) C57BL/6J (B6; n=10 M & n=5 F), +/+ (n=5 M & n=8 F), +/twgy (n=18 M & n=13 F) and twgy/twgy (n=36 M & n=39 F) mice are shown in (D). A treadmill exercise test run on +/+ (n=3 M & n=3 F), +/twgy (n=16 M & n=4 F) and twgy/twgy (n=6 M & n=9 F) mice showed impaired responses in homozygous twgy/twgy mice in time (in min; E), distance (in m; F), blood glucose (G) and lactate (H) at exhaustion. Means ± SD and individual data points are plotted with significant differences indicated (* = p < 0.05, ** = p < 0.01, *** = p < 0.001, **** = p < 0.0001).

When mice of the B/Int-x (N12F2) line were housed with access to a running wheel, only those mice with the homozygous twgy/twgy genotype showed the behavioral pattern identified in the original two lines (Fig. 7 A and Fig. 1 A-D). Thus, this pattern, consisting of decreased wheel-running behavior (Fig. 7 B) with a characteristic non-consolidated, messy pattern of expression, an increase in activity in the light phase (Fig. 7 C) and increased number of shorter, distinct bouts of activity, that is indicative of an overall excessive fatigability and sleep/wake disruption, shows a clear recessive mode of inheritance.

**Fig. 7.**
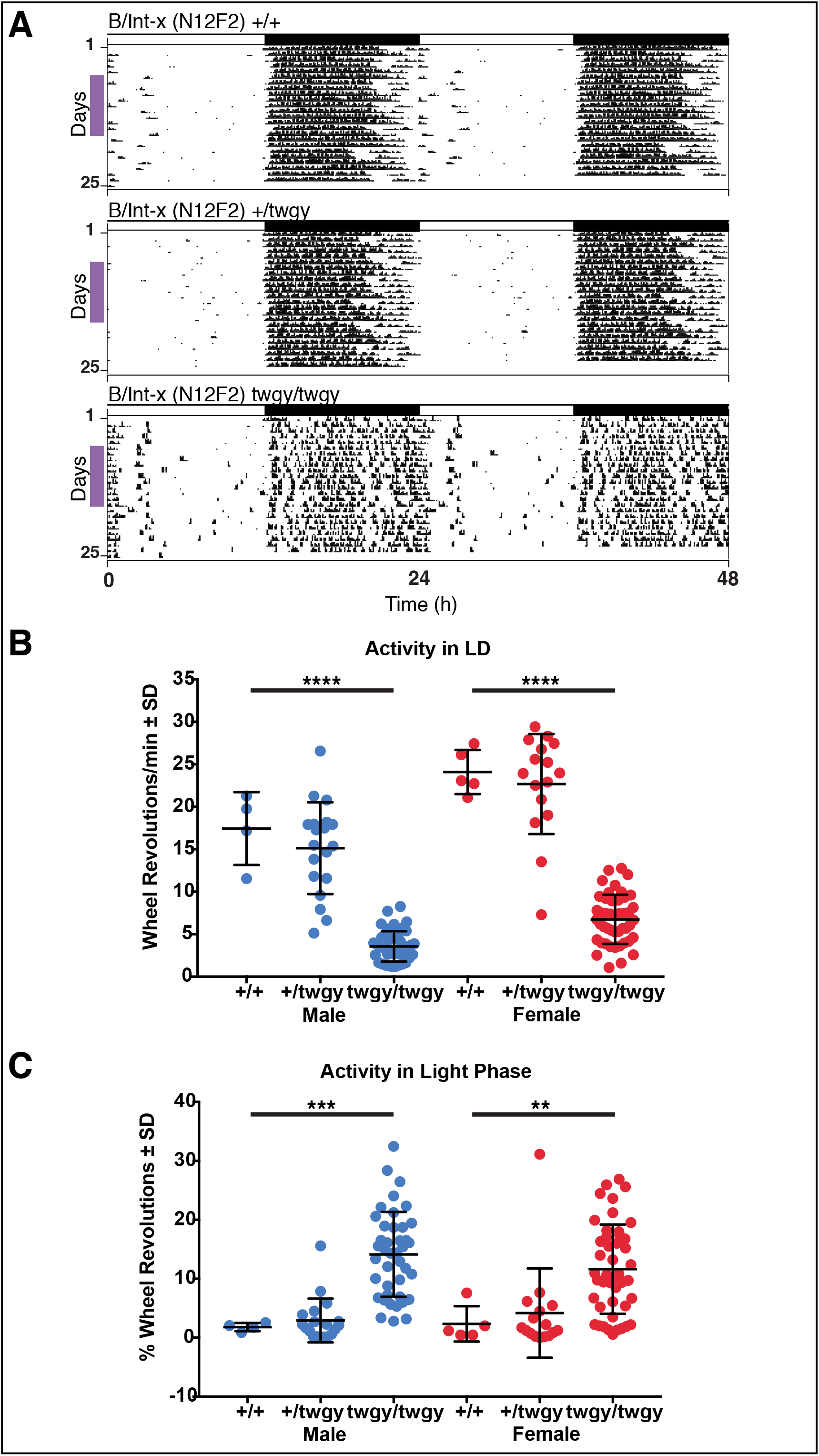
*Slc2a4^twgy/twgy^* mice show abnormal, low-amplitude circadian patterns of voluntary wheel-running behavior interrupted with frequent rest bouts. (A) Representative actograms of B/Int-x (N12F2) mice of all three genotypes (+/+ - top, +/twgy - middle, twgy/twgy - bottom) recorded for 3 weeks under a full 12:12 LD cycle (shown in bar across the top). Plotting conventions are like those for Fig 1A. Colored vertical bars on the left side of the actograms indicate days used for quantification of wheel-running behavior. Average number of wheel revolutions per min over 24h (B), and % of daily activity occurring in the light phase (C) for mice in the +/+ (n=4 M & n=5 F), +/twgy (n=19 M & n=16 F), and twgy/twgy (n=41 M & n=45 F) groups are plotted. Means ± SD and individual data points are plotted with significant differences indicated (** = p < 0.01, *** = p < 0.001, **** = p < 0.0001).

## SUMMARY AND DISCUSSION

Excessive fatigue that defies simple explanation affects large subsections of the population at one time or another, and results in diminished daily functioning and quality of life (1, 3, 4). It can be a symptom or side effect of varied diseases and disorders and their treatments, but can also be manifest independent of other comorbid diagnoses or etiologies (51-53). For those persons suffering from this debilitating disorder, and their loved ones, effective treatments would be most welcome, but in order for new treatments to be identified and developed, informative animal models of chronic fatigue are crucial. We carried out a forward genetic screen in order to identify mouse lines showing behavioral characteristic of chronic fatigue. We describe the identification of a novel mutation in the *Slc2a4* gene encoding GLUT4 that results in a distinct daily voluntary wheel-running pattern that is consistent with chronic fatigue. Affected mice homozygous for the *twiggy* mutation of *Slc2a4* show a behavioral pattern consisting of low, non-consolidated and messy wheel-running activity, with increased short bursts of activity interrupting the rest phase. These mice sleep during times of the day that normal nocturnal wildtype mice would never sleep (i.e., during the first half of the active phase). This pattern of behavior is consistent with increased fatigability, and suggests that impaired functioning of the insulin-responsive glucose transporter (GLUT4), or other impairments in glucose regulation and homeostasis may play a role in the underlying pathophysiology of chronic fatigue.

Additional phenotypic characteristics observed in *Slc2a4^twgy/twgy^* homozygous mice are consistent with a non-functioning GLUT4 transporter (46). These characteristics include a low body weight, impaired glucose tolerance response with a normal baseline fasting blood glucose level, increased heart mass consistent with cardiac hypertrophy, decreased exercise tolerance, and increased fatiguing response with abnormal blood glucose and lactate levels following exercise. Surprisingly, although these phenotypic characteristics largely match those described for the GLUT4 null mouse (46-49, 54-56), the protein levels in *Slc2a4^twgy/twgy^* homozygous mice were found to be elevated, suggestive of protein accumulation due to impairments in turnover or degradation of the protein, or compensatory upregulation. It is possible that the mutation results in a malformation of GLUT4 that interferes with its translocation or fusion to the plasma membrane, or that it impairs its function as a glucose transporter. Either of these possibilities may result in a compensatory over production of the protein. The premature stop introduced into exon 10 of the gene would likely result in the truncation of the protein and absence of the carboxy terminus. There are several important regulatory elements located in this region of the protein that play a role in its subcellular localization and transmission to the plasma membrane in response to circulating insulin, dietary components and metabolites, or exercise-related signaling pathways (57-62). It remains to be determined whether the phenotypes resulting from the presumably truncated form of *Slc2a4* are the result of impaired functioning of the glucose transporter itself, or from impairments in its vesicular localization and trafficking to the membrane in response to insulin or other signaling cascades.

The constellation of phenotypes described for mice in the *twiggy* lines, including low body weight, abnormal voluntary wheel running, and physiological alterations such as enlarged hearts and impaired glucose tolerance responses, all show a clear recessive heritance pattern. For virtually all phenotypes measured, heterozygous *Slc2a4^+/twgy^* mice were indistinguishable from their wildtype littermates and/or wildtype C57BL/6J controls. This is in sharp contrast to the GLUT4 knockout heterozygous mouse that exhibits phenotypes consistent with diabetes (63-66). It has been suggested that this diabetic phenotype, which is manifest in only a subpopulation of the heterozygous knockouts and appears to vary with age, could be an artifact of genetic background; however, this remains an open issue (64). In addition, GLUT4 protein levels are diminished in heterozygous GLUT4 knockout mice (63-66). In contrast, protein levels of GLUT4 were unchanged in the muscle of *Slc2a4^+/twgy^* mice. They were not diminished as in the heterozygous knockout, absent as in the homozygous knockout, nor elevated as in the *Slc2a4^twgy/twgy^* mice. This finding reinforces the idea that this mutation resulting in a premature stop in exon 10 of the gene does not simply knock out GLUT4, and therefore it represents a novel model that can be used to further characterize GLUT4 functioning.

The GLUT4 glucose transporter was historically considered a peripheral-acting glucose channel, with expression thought to be limited to muscle, adipose tissues, and heart (67, 68). Recent studies have shown that it is, in fact, expressed more widely and can also be found in distinct neuronal populations throughout the brain in mice (69). This more varied expression pattern and anatomical localization suggests that GLUT4 functions are more complex than simply a shunt for glucose in peripheral tissues. For example, a very recent study shows that GLUT4 functions to maintain neuronal, and specifically synaptic, energy requirements and is essential for maintaining extended synaptic transmission (70). The use of conditional genetic tools has also shed more light on the diverse functions of GLUT4. Interfering with insulin signaling specifically in all tissues expressing GLUT4 results in a diabetic phenotype that is due, at least in part, to impaired insulin signaling in neuronal populations (71). It is interesting to note that these mice showed decreased body weight, but did not show any alterations in activity amounts or patterns (71). Mice with GLUT4 removed selectively from the brain show that neuronal GLUT4 is critical for assessing and reacting to changes in blood glucose levels (72); however, it is possible that some of these effects are non-specific *Cre*-driver related since Nestin-*Cre* is expressed peripherally and mice carrying this transgene have been shown to have metabolic phenotypes (73). Work has been initiated to try to identify the function of distinct subgroups of GLUT4 neurons using methods to isolate and/or selectively ablate those expressing this gene (74, 75). The removal of hypothalamic neurons expressing GLUT4 resulted in metabolic changes that are consistent with impaired energy utilization and homeostatic processes, but did not result in locomotor deficits (75). These mice ate less and lost weight, had altered oxygen consumption and energy expenditure, and increased fasting blood glucose and aberrant glucose tolerance responses, but the amount and daily pattern of activity remained unchanged. These mice did, however, show a decreased activity response to an overnight fast, and did not increase activity levels as much as unablated controls (75). Clearly, the use of these genetic tools to conditionally manipulate the expression of GLUT4 in anatomically and temporally controlled ways will be critical in fully characterizing the function of this glucose transporter. This should allow for the determination of exactly how GLUT4 removal or altered function results in the seemingly varied array of phenotypic outcomes described in the literature to date.

The identification of a novel mutation in the *Slc2a4* gene resulting in a constellation of abnormal behavioral characteristics consistent with chronic fatigue was somewhat surprising, but not necessarily unexpected. It seems logical that impaired glucose processing and homeostasis, with a concomitant altered tolerance of exercise, would result in aberrant patterns of voluntary wheel-running activity as well. Voluntary wheel running is increased in mice overexpressing GLUT4 in muscle (76), but GLUT4 expression levels did not differ between active and inactive mouse strains (77). It is likely, however, that impaired glucose transport and cellular utilization would have some effect on voluntary sustained activity. Switching from predominantly glucose to fatty acid fuel utilization in muscle, either by transgenic or pharmacologic manipulation of the peroxisome proliferator-activated receptor delta (PPARδ) pathway, increases exercise endurance in mice (78-80). Whether or not this increase in exercise capacity translates into an increase in voluntary activity remains to be fully elucidated. For instance, mice deficient in both cryptochrome 1 and 2 (Cry1/2 dKO) ran faster and for longer distances than wildtype or single knockout controls on a treadmill task, presumably through de-repression of PPARδ activity; however, these mice showed decreased daily voluntary wheel-running behavior (81). It remains to be determined whether or not treatment of *Slc2a4^twgy/twgy^* mice with exercise mimetics would normalize some of the behavioral abnormalities in wheel running that we observed (82). This may prove informative to the investigation of how these novel pharmacological treatments may apply to patients with chronic fatigue, for whom voluntary exercise programs have been less than successful.

Fatigue, and alterations in the activity of daily living are well-described side effects and symptoms of both type I and type II diabetes (83-85). Like fatigue in other patient populations, fatigue in diabetes is associated with physiological, psychological and lifestyle variables, but may be exacerbated by fluctuations in blood glucose levels and homeostasis. There is also some evidence suggestive of abnormal metabolic responses in chronically fatigued patients, with abnormalities in energy and sugar metabolism comprising one of the three main categories of pathways identified (86). The identification of abnormalities in metabolic pathways in chronic fatigue, both as candidates for geared treatment options, and as potential biological markers that can be used to differentially diagnose various subtypes of the disorder, have received an increase in scientific focus recently (87, 88).

Using an unbiased, forward genetic ENU mutagenesis screen for chronic fatigue, we identified mice with a mutated form of the GLUT4 transporter; however, we are by no means suggesting that mutations in GLUT4, or impaired glucose transport underlie all manifestations of fatigue, although it may play a role in at least a subset of cases. Rather, it is likely that chronic fatigue represents a persistent altered homeostatic state in response to a variety of triggers and physiological adaptations (87). It is hoped that additional screening of mutagenized mouse lines for abnormalities in the expression of voluntary wheel-running behavior will yield additional candidate genes and pathways that are implemented in this complex disorder. In the mean time, studies to determine the effectiveness of treatment strategies geared towards normalizing glucose processing and transport in the *Slc2a4^twiggy^* mice and other mouse models of chronic fatigue may hold promise. In addition, assessing glucose tolerance in patients for whom fatigue remains an unexplained symptom may be warranted.

## MATERIALS AND METHODS

### Animals and Housing

All mice were group housed in standard mouse cages under a 12:12 LD cycle (lights on at 6:00 AM) with water and regular mouse chow (2018 Teklad Global 18% Protein Rodent Diet, Envigo, Madison, WI, USA) available *ad libitum*, unless otherwise stated. Mice were produced in our colony from strategic crosses, or were ordered from an in-house breeding core facility (Mouse Breeding Core, Wakeland lab, UT Southwestern Medical Center, Dallas, TX, USA), which orders its breeding mice from Jackson Labs (Bar Harbor, ME, USA). Some wildtype C57BL/6J mice were also ordered from Jackson Labs (Stock Number 000664) directly. The generation of the original ENU mutagenized screening population and body weight screen were done at Northwestern University, and all behavioral and subsequent phenotypic characterizations were performed at the University of Texas Southwestern Medical Center. The Institutional Animal Care and Use Committee (IACUC) of Northwestern University and the University of Texas Southwestern Medical Center approved all animal procedures (ACUC# 2003-0034, and APNs 2009-0054, 2015-100925 and 2015-101140, respectively). The Slc2a4^*twiggy*^ line has been donated to Jackson Labs (C57BL/6J-*Slc2a4^twiggy^*/J, Stock Number 029699).

### ENU mutagenesis and Phenotypic Screening

The generation of N-Ethyl-N-nitrosourea (ENU; Sigma; catalog #N3385) mutagenized mice and their progeny for initial high-throughput screening was described previously (45). Briefly, male C57BL/6J mice (~6 weeks of age) were injected with 250 mg/kg of ENU. Following a recovery period of ~6 weeks, these mice were crossed with wildtype C57BL/6J females to produce generation 1 (G1) males which were again crossed to wildtype C57BL/6J females to produce G2 females. These G2 females were backcrossed to their G1 fathers to produce G3 mice of both sexes used for phenotype screening.

More advanced generations of mice, further removed from the original G0 founders, were used for the more in depth behavioral phenotypic screening. Progeny of selected lines of mice that were at least 8 weeks old were individually housed in standard polycarbonate mouse cages (Fischer Scientific; catalog # 01-288-1B and 01-288-21) containing a 4.75” diameter stainless steel running wheel placed inside isolation cabinets containing 12 to 24 cages each, as described previously (19). Temperature and humidity were monitored throughout, and the mice were initially recorded under a 12:12 LD cycle (green LEDs, ~100 lux at the level of the cage floor). Following the initial 3 weeks of testing under 12:12 LD, selected individual progeny from specific crosses were tested under constant darkness (DD) or constant light (LL) conditions. Wheel-running behavior was recorded using ClockLab Data Acquisition Software (Actimetrics Inc., Wilmette, IL, USA). Wheel-running data, specifically total daily amount, number of daily bouts, and percentage of activity occurring in the light during LD recording, as well as total daily amount of activity, free-running period, and amplitude of the rhythm recorded in DD, were quantified using ClockLab Analysis Software (Actimetrics Inc., Wilmette, IL, USA). For all data quantification, recording days 6-15 under LD and 31-40 under DD were used. All wheel-running cages, food and water bottles were changed every 21 days.

### Behavioral Time-Lapsed Photography and Video Recording

I-x (N2F14 and N2F15) and C57BL/6J wildtype mice were housed in individual running-wheel cages as described above. Using a Sony Handycam DCR-HC62 camera with an LED IR light source, photographs were taken at 1 min intervals beginning at lights off (or 18:00) for 15h. The resulting pictures were scored from 1 to 9 for the behavioral state or cage location depicted in each. The scores used were: 1) drinking, 2) eating, 3) near the food hopper, 4) on the wheel running, 5) near the wheel awake, 6) end of the cage awake, 7) end of the cage resting, 8) near the wheel asleep, and 9) end of the cage asleep. These scores were plotted in ethograms according to the time at which each picture was taken.

Video recording of 3 I-x (N2F21) mice, and mice from two different test crosses (T-x (N3F2) n=2 and T-x (N11F3) n=3), was done under the same IR light source described above using the same Sony Handycam camera with Sony Premium Mini DV Cassettes (DVM60 ME LP:90) from 20:00 to 21:30 (i.e., 2h to 3.5h into the dark phase). Wheel running was recorded throughout this period as described above. The resulting video was scored for behavioral arrest indicative of sleep by two researchers blind to mouse identity. Periods of immobility with lowered posture lasting longer than 40 consecutive seconds were scored as sleep (89).

### Body Weight, Fasting Blood Glucose, Health Assessment, and Tissue Collections

All mice produced in our colony were weighed at specific ages through development. At weaning, and at regular intervals thereafter, mice were visually inspected for general health and wellness. For the determination of blood glucose levels, mice were fasted for 4h during the middle of the light phase (a time when mice do not normally eat). Body weights were measured at the beginning and the end of the 4h fast, and blood glucose was determined from a small droplet of blood taken from the tip of the tail, using the Precision Xtra blood glucose meters and glucose test strips (Abbott Diabetes Care Inc., Almeda, CA, USA). At the end of experiments and phenotypic characterization, mice were euthanized with CO_2_ followed by cervical dislocation. In some cases, tissues (tailtips, spleens and hearts) were collected for weighing or DNA extraction followed by genotyping.

### Mapping

Affected mice from the I-x and B/Int-x lines were selected and crossed with wildtype C57BL/10J mice (Stock Number 000665, Jackson Labs, Bar Harbor, ME, USA) to produce mapping (M-x) F1 mice that were intercrossed to produce F2 progeny. Additional crosses of wildtype C57BL/6J and C57BL/10J were also set up to produce control (WTB6B10) F1 and F2 mice. All mice produced were individually housed in running-wheel cages after 8 weeks of age and their behavior was recorded in LD (for 3 weeks) and DD (for 3 weeks) as described above. Following behavioral testing, mice were euthanized and spleens were collected, frozen onto dry ice, and then stored at - 80°C. Spleen DNA from 74 selected mice (half presumptive homozygous mutants and half presumptive wildtypes or heterozygotes) was extracted using phenol chloroform genomic DNA extraction method. The final DNA pellet was suspended in RNAse free H2O. Concentrations were determined with a NanoDrop spectrophotometer, and DNA was genotyped using Taqman probes on Fluidigm platform using the 96.96 chip according to manufacturer instructions as described previously (90). QTL linkage analysis was performed using R/QTL as described previously (90, 91). Genotyping failed for one spleen sample, and these data were subsequently removed from further analyses.

### Whole Exome Sequencing

Test crosses (T-x) of affected, presumably homozygous B/Int-x (N11F2) were set up to produce homozygous T-x (N11F3) progeny. Following phenotype recording on wheels, mice were euthanized and spleens were collected from 16 representative T-x (N11F3) individuals and 16 representative I-x (N2F20) mice. Spleens were quickly frozen on dry ice and were then stored at -80°C until DNA was extracted using the Gentra Puregene Mouse Tail/Tissue Kit (Catalog number 158906 - Cell Lysis Solution, 158918 - Puregene Proteinase K, 158922 - RNase A Solution, and 158910 - Protein Precipitation Solution; Qiagen, Germantown, MD, USA) according to manufacturer instructions. DNA was dissolved in sterile RNAse free H2O and concentrations were determined using a NanoDrop spectrophotometer (ND-2000, ThermoFisher Scientific, Waltham, MA, USA) and Qubit 2.0 fluorometric quantitation (Q32866, ThermoFisher Scientific, Waltham, MA, USA). DNA samples for the 16 individuals of each of the two lines were pooled into two samples that were used for whole exome sequencing.

Whole-exome capture, sequencing, and analysis were performed by the UT Southwestern McDermott Center Next-Generation Sequencing and Bioinformatics Core. After initial DNA quality assessment on a 1.8% agarose gel and concentration quantification using a Qubit 3.0 Fluorometer (Invitrogen, Carlsbad, CA USA), fragment libraries were prepared using SureSelectXT2 HSQ Reagent Kit (Agilent Technologies, Inc., Santa Clara, CA, USA). Samples were sheared on the Covaris S-2 sonicator (Covaris, Woburn, MA, USA) and end-repaired, after which the 3’ ends were adenylated and barcoded with pre-capture indexing adapters. Following amplification and purification, the sizes of the fragment libraries were assessed on the Agilent 2100 BioAnalyzer (Agilent Technologies, Inc., Santa Clara, CA, USA) and concentrations were once again determined by Qubit fluorometric quantitation. Samples were pooled in equimolar amounts and captured with the SureSelectXT2 Target Enrichment System for Mouse (Agilent Technologies, Inc., Santa Clara, CA, USA), followed by a final Qubit concentration quantification, and were run on the Illumina NextSeq 500 platform (Illumina Inc., San Diego, CA, USA) using 150PE SBS v2 chemistry. Sequence reads were mapped to the Genome Reference Consortium GRCm38 reference genomic sequence for C57BL/6J using BWA (92). Both samples resulted in mean coverage greater than 120X. Variants were called using the Genome Analysis ToolKit (GATK) (93), and annotated using snpEff (94). Reads were visualized and plotted using the Broad Institute Integrative Genomics Viewer (IGV) to detect homozygous mutations present in both samples.

### Genotyping

NaOH-extracted DNA from tailtip samples taken at weaning from progeny of selected crosses was genotyped for the presence of the *twiggy* mutation with real-time PCR to detect the SNP using the following primers: Slc2a4 Forward (ccagagaccacctacatggc) Slc2a4 WTReverse (GTCGGCATGGGTTTCCAGaAT) and Slc2a4 MutantReverse (GTCGGCATGGGTTTCCAGaAa). PCR reactions were done in a 10 µl volume using ~10 ng genomic DNA template with KAPA SYBR Fast ABI Prism 2X qPCR Master Mix (KK4617; KAPA Biosystems, Wilmington, MA, USA), on an ABI 7900HT Fast Real-Time PCR Machine (Applied Biosystem, Inc., Foster City, CA, USA) using MicroAmp 384-well Optical Reaction Plates (Applied Biosystems, Inc., Foster City, CA, USA) with the following thermo-cycling conditions: 95°C for 3 min followed by 40 cycles of 95°C for 3 sec and 62°C for 20 sec.

### Glucose Tolerance Test

Progeny of select crosses were fasted for 4h during the middle of the light phase (a time when mice do not normally eat). Body weights were measured at the beginning and the end of the 4h fast, and initial blood glucose was determined from a small droplet of blood taken from the tip of the tail using the Precision Xtra blood glucose meters and glucose test strips (Abbott Diabetes Care Inc., Almeda, CA, USA). Mice were then given an I.P. injection of 10% sterile glucose solution (1 g/kg; Sigma-Aldrich, St. Louis, MO, USA), and blood glucose values were measured again at 15, 30, 45, 60 and 120 min post injection.

### Endurance Treadmill Test

To test exercise endurance capacity, mice of both sexes from selected crosses were run on a progressive treadmill running protocol. The mice were acclimatized to the motorized treadmill (Animal Treadmill Exer-3/6, Columbus Instruments, Columbus, OH, USA) for two days prior to the endurance test. Day 1 of acclimatization lasted 20 min consisting of 5 min on the treadmill at rest (0 m/min), 5 min at 8 m/min, 5 min at 10 m/min, followed by 5 min of rest (0 m/min). Day 2 also lasted 20 min and consisted of 5 min on the treadmill at rest (0 m/min), 5 min at 10 m/min, 5 min at 12 m/min, followed by 5 min of rest (0 m/min). On day 3, mice performed the endurance test to exhaustion, defined as when the mouse spent more than 5 sec on the shock grid (0.1 mA x163 V and 1 Hz) without re-engaging with the treadmill, as described previously (95). The mice ran at a speed of 10 m/min for the first 40 min, then the rate was increased at 1 m/min every 10 min until the speed reached 13 m/min, at which point the speed was increased at a rate of 1 m/min every 5 min until exhaustion. Mice were then removed from the treadmill and blood glucose and lactate were immediately measured from tail blood using hand-held analyzers for glucose (Bayer Contour Blood Glucose Meter, Ascensia Diabetes Care US, Inc., Parsippany, NJ, USA) and lactate (Nova Biomedical Lactate Plus Meter, Waltham, MA, USA). Mice were euthanized by decapitation, and muscles were rapidly dissected and frozen on dry ice. All tissues were stored at -80°C until used.

### Western Blotting

Frozen gastrocnemius muscles collected from mice following the treadmill study were homogenized in ice-cold lysis buffer composed of T-PER^®^ buffer (#78510, ThermoFisher Scientific, Waltham, MA, USA), 1% (v/v) of protease inhibitors cocktail (P8340-5ML, Sigma-Aldrich, St. Louis, MO, USA), and phosphatase inhibitors cocktail 2 and 3 (P5726-5ML and P0044-5ML, Sigma-Aldrich, St. Louis, MO, USA). Samples were homogenized using TissueLyser II homogenizer (Qiagen Inc., Valencia, CA, USA). Homogenates were then solubilized by constant rotation for 1h at 4°C, and 10 min x 1,000 *g* at 4°C. Protein concentration of the supernatant was determined via the Pierce BCA protein assay kit (#23225, ThermoFisher Scientific, Waltham, MA, USA). Samples were then stored at -80°C until further analysis. Equal amounts of total protein (20 µg) per sample were diluted with appropriate volume of laemmle sample buffer (2X concentrated; 4% SDS, 10% 2-mercaptoethanol, 20% glycerol, 0.004% bromophenol blue and 0.125 M pH6.8 Tris-HCl), heated for 5 min at 95°C, separated via SDS-PAGE, and transferred to nitrocellulose (Trans-blot turbo, Bio-Rad, Hercules, CA, USA). Membranes were incubated with the appropriate primary (GLUT4 #2213 and β-Actin #3700, Cell Signaling Technologies, Danvers, MA, USA) and fluorescent secondary antibodies (IRDye 680 Goat anti-Mouse IgG, Li-Cor Bioscience, Lincoln, NE, USA). Protein band fluorescence was quantified via Li-Cor Odessy Image studio Version 4.0 (Li-Cor Bioscience, Lincoln, NE, USA). Individual values are relative to the mean of all sample values within the same membrane, and equal loading was confirmed and normalized to the immuno-reactivity of β-Actin.

### Data Analysis and Statistics

All data were analyzed and data plots were generated using Prism 6 (GraphPad Software, Inc., La Jolla, CA, USA). Statistical comparisons were made using ANOVA with post hoc analyses to compare groups. Tests with p < 0.05 were deemed statistically significant. Unless otherwise stated, all values plotted are presented as mean ± SD, with statistical results presented as * = p < 0.05, ** = p < 0.01, *** = p < 0.001 and **** = p < 0.0001.

## AUTHOR CONTRIBUTIONS

Experiments were designed by MHMdG, CMC, VK, JAM, and JST. Animal care and line maintenance was done by MHMdG. MHMdG, CMC, VK, JAM, and NIA performed the experiments and collected the data. Data analysis, interpretation and the generation of figures was done by MHMdG, CMC, VK, JAM and JST. The manuscript was written, reviewed and edited by MHMdG, CMC, VK, JAM and JST. JST directed and funded the research.

## Acknowledgements

### ACKNOWLEDGMENTS

Research was supported by the NIH (#U01 MH 61915) and the Howard Hughes Medical Institute (JST). Additional funding was received by CMC from the NIDDK (F32DK104659). We would like to thank Delali Bassowou, Kelly Bruckmann, Kelly Foster, Kyung-Inn Kim, Izabela Kornblum, Sammie McMurray, Chris Olker, and Dawn Olson for technical assistance. We thank Sandy Siepka and Martha Vitaterna for production of the ENU mutagenized mice and design of the initial phenotypic screen. JST is a co-founder of, a Scientific Advisory Board member of, and a paid consultant for Reset Therapeutics, Inc., a biotechnology company aimed at discovering small-molecule therapies that modulate circadian activity for a variety of disease indications. JST is an Investigator, and MHMdG. is a Research Specialist, in the Howard Hughes Medical Institute.

## REFERENCES

1 Jason LA, Evans M, Brown M, & Porter N (2010) What is fatigue? Pathological and nonpathological fatigue. PM R 2(5):327–331.

2 Fukuda K, et al. (1994) The chronic fatigue syndrome: A comprehensive approach to its defintion and study. Ann Intern Med 121:953–959.

3 Wessely S (1998) The epidemiology of chronic fatigue syndrome. Epidemiol Psichiatr Soc 7(1):10–24.

4 Ranjith G (2005) Epidemiology of chronic fatigue syndrome. Occup Med (Lond) 55(1):13–19.

5 Hardy SE & Studenski SA (2008) Fatigue predicts mortality in older adults. J Am Geriatr Soc 56(10):1910–1914.

6 Hardy SE & Studenski SA (2008) Fatigue and function over three years among older adults. J Gerontol A Biol Sci Med Sci 63(12):1389–1392.

7 Moreh E, Jacobs JM, & Stessman J (2010) Fatigue, function, and mortality in older adults. J Gerontol A Biol Sci Med Sci 65(8):887–895.

8 Maes M & Twisk FNM (2010) Chronic fatigue syndrome: Harvey and Wessely’s (bio)psychosocial model versus a bio(psychosocial) model based on inflammatory and oxidative and nitrosative stress pathways. BMC Medicine 8(35):1–13.

9 Harrington ME (2012) Neurobiological studies of fatigue. Prog Neurobiol 99(2):93–105.

10 Anisman H, Gibb J, & Hayley S (2008) Influence of continuous infusion of interleukin-1beta on depression-related processes in mice: corticosterone, circulating cytokines, brain monoamines, and cytokine mRNA expression. Psychopharmacology (Berl) 199(2):231–244.

11 Kudo T, Loh DH, Truong D, Wu Y, & Colwell CS (2011) Circadian dysfunction in a mouse model of Parkinson’s disease. Exp Neurol 232(1):66–75.

12 Kudo T, et al. (2011) Dysfunctions in circadian behavior and physiology in mouse models of Huntington’s disease. Exp Neurol 228(1):80–90.

13 Nakamura TJ, et al. (2011) Age-related decline in circadian output. J Neurosci 31(28):10201–10205.

14 Ottenweller JE, et al. (1998) Mouse Running Activity Is Lowered by Brucella abortus Treatment: A Potential Model to Study Chronic Fatigue. Physiol Behav 63(5):795–801.

15 Ray MA, Trammell RA, Verhulst S, Ran S, & Toth LA (2011) Development of a Mouse Model for Assessing Fatigue during Chemotherapy. Comp Medicine 61(2):119–130.

16 Bonsall DR, et al. (2015) Suppression of Locomotor Activity in Female C57Bl/6J Mice Treated with Interleukin-1beta: Investigating a Method for the Study of Fatigue in Laboratory Animals. PLoS One 10(10):e0140678.

17 Lightfoot JT (2011) Current understanding of the genetic basis for physical activity. J Nutr 141(3):526–530.

18 Shimomura K, et al. (2001) Genome-wide epistatic interaction analysis reveals complex genetic determinants of circadian behavior in mice. Genome Res 11(6):959–980.

19 Siepka SM & Takahashi JS (2005) Methods to record circadian rhythm wheel running activity in mice. Methods Enzymol 393:230–239.

20 Schur E, Afari N, Goldberg J, Buchwald D, & Sullivan PF (2007) Twin analyses of fatigue. Twin Res Hum Genet 10(5):729–733.

21 Landmark-Hoyvik H, et al. (2010) The genetics and epigenetics of fatigue. PM R 2(5):456–465.

22 Sprangers MA, et al. (2014) Biological pathways, candidate genes, and molecular markers associated with quality-of-life domains: an update. Qual Life Res 23(7):1997–2013.

23 Wang T, Yin J, Miller AH, & Xiao C (2017) A systematic review of the association between fatigue and genetic polymorphisms. Brain Behav Immun 62:230–244.

24 Narita M, et al. (2003) Association between serotonin transporter gene polymorphism and chronic fatigue syndrome. Biochem Biophys Res Commun 311(2):264–266.

25 Torpy DJ & Ho JT (2007) Corticosteroid-binding globulin gene polymorphisms: clinical implications and links to idiopathic chronic fatigue disorders. Clin Endocrinol (Oxf) 67(2):161–167.

26 Collado-Hidalgo A, Bower JE, Ganz PA, Irwin MR, & Cole SW (2008) Cytokine gene polymorphisms and fatigue in breast cancer survivors: early findings. Brain Behav Immun 22(8):1197–1200.

27 Lin E & Huang LC (2008) Identification of significant genes in genomics using Bayesian variable selection methods. Adv Appl Bioinform Chem 1:13–18.

28 Smith AK, et al. (2008) Genetic evaluation of the serotonergic system in chronic fatigue syndrome. Psychoneuroendocrinology 33(2):188–197.

29 Smith AK, Fang H, Whistler T, Unger ER, & Rajeevan MS (2011) Convergent genomic studies identify association of GRIK2 and NPAS2 with chronic fatigue syndrome. Neuropsychobiology 64(4):183–194.

30 Johnston S, Staines D, Klein A, & Marshall-Gradisnik S (2016) A targeted genome association study examining transient receptor potential ion channels, acetylcholine receptors, and adrenergic receptors in Chronic Fatigue Syndrome/Myalgic Encephalomyelitis. BMC Med Genet 17(1):79.

31 Schlauch KA, et al. (2016) Genome-wide association analysis identifies genetic variations in subjects with myalgic encephalomyelitis/chronic fatigue syndrome. Transl Psychiatry 6:e730.

32 Petersen HH, et al. (2006) Hyporesponsiveness to glucocorticoids in mice genetically deficient for the corticosteroid binding globulin. Mol Cell Biol 26(19):7236–7245.

33 Golumbek PT, Keeling RM, & Connolly AM (2007) RAG2 gene knockout in mice causes fatigue. Muscle Nerve 36(4):471–476.

34 Krzyszton CP, et al. (2008) Exacerbated fatigue and motor deficits in interleukin-10-deficient mice after peripheral immune stimulation. Am J Physiol Regul Integr Comp Physiol 295(4):R1109–1114.

35 Justice MJ, Noveroske JK, Weber JS, Zheng B, & Bradley A (1999) Mouse ENU Mutagenesis. Human Mol Genetics 8(10):1955–1963.

36 Soewarto D, et al. (2000) The large-scale Munich ENU-mouse-mutagenesis screen. Mamm Genome 11(7):507–510.

37 Keays DA & Nolan PM (2003) N-ethyl-N-nitrosourea mouse mutants in the dissection of behavioural and psychiatric disorders. European Journal of Pharmacology 480(1–3):205–217.

38 Clark AT, et al. (2004) Implementing large-scale ENU mutagenesis screens in North America. Genetica 122(1):51–64.

39 Goldowitz D, et al. (2004) Large-scale mutagenesis of the mouse to understand the genetic bases of nervous system structure and function. Brain Res Mol Brain Res 132(2):105–115.

40 Cordes SP (2005) N-ethyl-N-nitrosourea mutagenesis: boarding the mouse mutant express. Microbiol Mol Biol Rev 69(3):426–439.

41 Vitaterna MH, Pinto LH, & Takahashi JS (2006) Large-scale mutagenesis and phenotypic screens for the nervous system and behavior in mice. Trends Neurosci 29(4):233–240.

42 Kumar V, et al. (2011) Second-generation high-throughput forward genetic screen in mice to isolate subtle behavioral mutants. Proc Natl Acad Sci U S A 108(3):15557–15564.

43 Moresco EM, Li X, & Beutler B (2013) Going forward with genetics: recent technological advances and forward genetics in mice. Am J Pathol 182(5):1462–1473.

44 Takahashi JS, Shimomura K, & Kumar V (2008) Searching for genes underlying behavior: lessons from circadian rhythms. Science 322(5903):909–912.

45 Siepka SM & Takahashi JS (2005) Forward genetic screens to identify circadian rhythm mutants in mice. Methods Enzymol 393:219–229.

46 Katz EB, Stenbit AE, Hatton K, DePinho R, & Charron MJ (1995) Cardiac and adipose tissue abnormalities but not diabetes in mice deficient in GLUT4. Nature 377(6545):151–155.

47 Gorselink M, et al. (2002) Increased muscle fatigability in GLUT-4-deficient mice. Am J Physiol Endocrinol Metab 282(2):E348–354.

48 Simoes MV, et al. (2004) Delayed response of insulin-stimulated fluorine-18 deoxyglucose uptake in glucose transporter-4-null mice hearts. J Am Coll Cardiol 43(9):1690–1697.

49 Fueger PT, et al. (2007) Glucose kinetics and exercise tolerance in mice lacking the GLUT4 glucose transporter. J Physiol 582(Pt 2):801–812.

50 Fam BC, et al. (2012) Normal muscle glucose uptake in mice deficient in muscle GLUT4. J Endocrinol 214(3):313–327.

51 Morris G, Berk M, Walder K, & Maes M (2015) The Putative Role of Viruses, Bacteria, and Chronic Fungal Biotoxin Exposure in the Genesis of Intractable Fatigue Accompanied by Cognitive and Physical Disability. Mol Neurobiol 53(4):2550–2571.

52 Saligan LN, et al. (2015) The biology of cancer-related fatigue: a review of the literature. Support Care Cancer 23(8):2461–2478.

53 Skoie IM, Ternowitz T, Jonsson G, Norheim K, & Omdal R (2015) Fatigue in psoriasis: a phenomenon to be explored. Br J Dermatol 172(5):1196–1203.

54 Tsao TS, et al. (1997) Muscle-specific transgenic complementation of GLUT4-deficient mice. Effects on glucose but not lipid metabolism. J Clin Invest 100(3):671–677.

55 Stenbit AE, et al. (2000) Preservation of glucose metabolism in hypertrophic GLUT4-null hearts. Am J Physiol Heart Circ Physiol 279(1):H313–318.

56 Jiang H, Li J, Katz EB, & Charron MJ (2001) GLUT4 ablation in mice results in redistribution of IRAP to the plasma membrane. Biochem Biophys Res Commun 284(2):519–525.

57 Huang S & Czech MP (2007) The GLUT4 glucose transporter. Cell Metab 5(4):237–252.

58 Sadler JB, Bryant NJ, Gould GW, & Welburn CR (2013) Posttranslational modifications of GLUT4 affect its subcellular localization and translocation. Int J Mol Sci 14(5):9963–9978.

59 Govers R (2014) Cellular regulation of glucose uptake by glucose transporter GLUT4. Adv Clin Chem 66:173–240.

60 Alvim RO, Cheuhen MR, Machado SR, Sousa AG, & Santos PC (2015) General aspects of muscle glucose uptake. An Acad Bras Cienc 87(1):351–368.

61 Gannon NP, Conn CA, & Vaughan RA (2015) Dietary stimulators of GLUT4 expression and translocation in skeletal muscle: a mini-review. Mol Nutr Food Res 59(1):48–64.

62 Sylow L, Kleinert M, Richter EA, & Jensen TE (2017) Exercise-stimulated glucose uptake - regulation and implications for glycaemic control. Nat Rev Endocrinol 13(3):133–148.

63 Rossetti L, et al. (1997) Peripheral but not hepatic insulin resistance in mice with one disrupted allele of the glucose transporter type 4 (GLUT4) gene. J Clin Invest 100(7):1831–1839.

64 Stenbit AE, et al. (1997) GLUT4 heterozygous knockout mice develop muscle insulin resistance and diabetes. Nat Med 3(10):1096–1101.

65 Tsao TS, et al. (1999) Prevention of insulin resistance and diabetes in mice heterozygous for GLUT4 ablation by transgenic complementation of GLUT4 in skeletal muscle. Diabetes 48(4):775–782.

66 Li J, Houseknecht KL, Stenbit AE, Katz EB, & Charron MJ (2000) Reduced glucose uptake precedes insulin signaling defects in adipocytes from heterozygous GLUT4 knockout mice. FASEB J 14(9):1117–1125.

67 Charron MJ, Brosius FC, Alper SL, & Lodish HF (1989) A glucose transport protein expressed predominately in insulin-responsive tissues. Proc Natl Acad Sci USA 86:2535–2539.

68 Liu ML, et al. (1992) Expression and regulation of the human GLUT4/muscle-fat facilitative glucose transporter gene in transgenic mice. J Biol Chem 267(17):11673–11676.

69 Choeiri C, Staines W, & Messier C (2002) Immunohistochemical localization and quantification of glucose transporters in the mouse brain. Neuroscience 111(1):19–34.

70 Ashrafi G, Wu Z, Farrell RJ, & Ryan TA (2017) GLUT4 Mobilization Supports Energetic Demands of Active Synapses. Neuron 93(3):606–615 e603.

71 Lin HV, et al. (2011) Diabetes in mice with selective impairment of insulin action in Glut4-expressing tissues. Diabetes 60(3):700–709.

72 Reno CM, et al. (2017) Brain GLUT4 Knockout Mice Have Impaired Glucose Tolerance, Decreased Insulin Sensitivity, and Impaired Hypoglycemic Counterregulation. Diabetes 66(3):587–597.

73 Harno E, Cottrell EC, & White A (2013) Metabolic pitfalls of CNS Cre-based technology. Cell Metab 18(1):21–28.

74 Ren H, et al. (2014) Glut4 expression defines an insulin-sensitive hypothalamic neuronal population. Mol Metab 3(4):452–459.

75 Ren H, Lu TY, McGraw TE, & Accili D (2015) Anorexia and Impaired Glucose Metabolism in Mice With Hypothalamic Ablation of Glut4 Neurons. Diabetes 64:405–417.

76 Tsao TS, et al. (2001) Metabolic adaptations in skeletal muscle overexpressing GLUT4: effects on muscle and physical activity. FASEB J 15(6):958–969.

77 Dawes M, et al. (2014) Differential gene expression in high- and low-active inbred mice. BioMed research international 2014:361048.

78 Wang YX, et al. (2004) Regulation of muscle fiber type and running endurance by PPARdelta. PLoS Biol 2(10):e294.

79 Narkar VA, et al. (2008) AMPK and PPARdelta agonists are exercise mimetics. Cell 134(3):405–415.

80 Fan W, et al. (2017) PPARdelta Promotes Running Endurance by Preserving Glucose. Cell Metab 25(5):1186–1193 e1184.

81 Jordan SD, et al. (2017) CRY1/2 Selectively Repress PPARdelta and Limit Exercise Capacity. Cell Metab 26(1):243–255 e246.

82 Fan W & Evans RM (2017) Exercise Mimetics: Impact on Health and Performance. Cell Metab 25(2):242–247.

83 Fritschi C & Quinn L (2010) Fatigue in patients with diabetes: a review. J Psychosom Res 69(1):33–41.

84 Lasselin J, et al. (2012) Fatigue and cognitive symptoms in patients with diabetes: relationship with disease phenotype and insulin treatment. Psychoneuroendocrinology 37(9):1468–1478.

85 Moulton CD, Pickup JC, & Ismail K (2015) The link between depression and diabetes: the search for shared mechanisms. Lancet Diabetes Endocrinol 3(6):461–471.

86 Germain A, Ruppert D, Levine SM, & Hanson MR (2017) Metabolic profiling of a myalgic encephalomyelitis/chronic fatigue syndrome discovery cohort reveals disturbances in fatty acid and lipid metabolism. Mol Biosyst 13(2):371–379.

87 Klimas NG, Broderick G, & Fletcher MA (2012) Biomarkers for chronic fatigue. Brain Behav Immun 26(8):1202–1210.

88 Maxmen A (2017) Biological underpinnings of chronic fatigue syndrome begin to emerge. Nature 543(7647):602.

89 Pack AI, et al. (2007) Novel method for high-throughput phenotyping of sleep in mice. Physiol Genomics 28(2):232–238.

90 Kumar V, et al. (2013) C57BL/6N mutation in cytoplasmic FMRP interacting protein 2 regulates cocaine response. Science 342(6165):1508–1512.

91 Broman KW, Wu H, Sen S, & Churchill GA (2003) R/qtl: QTL mapping in experimental crosses. Bioinformatics 19(7):889–890.

92 Li H & Durbin R (2009) Fast and accurate short read alignment with Burrows-Wheeler transform. Bioinformatics 25(14):1754–1760.

93 McKenna A, et al. (2010) The Genome Analysis Toolkit: a MapReduce framework for analyzing next-generation DNA sequencing data. Genome Res 20(9):1297–1303.

94 Cingolani P, et al. (2012) A program for annotating and predicting the effects of single nucleotide polymorphisms, SnpEff: SNPs in the genome of Drosophila melanogaster strain w1118; iso-2; iso-3. Fly (Austin) 6(2):80–92.

95 Fujikawa T, et al. (2016) SF-1 expression in the hypothalamus is required for beneficial metabolic effects of exercise. Elife 5.

